# Dynamic Price and Breeder’s equations for variable environments

**DOI:** 10.1101/762658

**Authors:** Tim Coulson, Tomos Potter, Anja Felmy

## Abstract

The Breeder’s and Price equations are static models of evolution. Recent work has demonstrated how components of these equations, such as selection differentials and heritabilities, can be calculated for each time-step from dynamic, evolutionarily-explicit, structured population models. These dynamic models consist of functions that describe how environmental factors impact associations of genotypes and phenotypic traits with survival, reproduction, development, and non-genetic inheritance. Because evolutionarily explicit versions of these structured models can include feedbacks and can capture all the routes via which environmental variation can impact evolution, they i) provide a more powerful predictive tool for modelling eco-evolution in natural settings than existing approaches, and ii) reveal how key evolutionary parameters incorporated into static models change with time as evolution, or environmental change, proceeds.

## INTRODUCTION

Eco-evolutionary dynamics occur when selection on heritable phenotypic traits and the population growth rate within a focal species change together (Hairston Jr et al. 2005). Eco-evolutionary feedbacks then arise when a change in the population growth rate alters the strength of selection within a species, and vice versa (Patel et al. 2018). By using dynamic, evolutionarily-explicit, structured models (Coulson et al. 2017) to examine the dynamics of both population growth and terms in the Breeder’s and Price equations (Coulson et al. 2010, Childs et al. 2016), we show how eco-evolutionary dynamics and feedbacks influence evolution, phenotypic trait dynamics, and population growth. Our approach is based on understanding the interplay of fitness functions with functions that describe development, and genetic and non-genetic inheritance (see also (Wallace et al. 2013)). Fitness functions define how the association between phenotypic trait values and individual absolute fitness (or survival and reproductive components of fitness) varies with environmental drivers; development functions characterise the phenotypic development of surviving individuals over a time-step as a function of their environment; inheritance functions capture both genetic inheritance and environmentally-induced, non-genetic inheritance (Childs et al. 2016, Coulson et al. 2017).

Adaptive evolution requires directional selection on a heritable phenotypic trait (Fisher 1930). The strength of selection is determined by the phenotypic variance and the association between the phenotypic trait and absolute fitness, and is measured with a selection differential (Price 1970, Lande and Arnold 1983). Selection alters the distribution of the trait within a population within a generation. For evolution to occur, this change must, at least partially, be reflected in the distributions of alleles that determine trait values within the population (Falconer 1960). Rules of genetic inheritance mean that allele frequencies among selected parents are expected to be observed in offspring, although processes such as mutation and genetic drift can lead to departures from this expectation (Crow and Kimura 1970). For eco-evolutionary dynamics and feedbacks to occur, it is necessary, but not sufficient, that the strength of selection is influenced by the population growth rate. The selection differential can be written as the covariance between phenotypic trait values and individual absolute fitness, divided by the mean individual absolute fitness (Heywood 2005). When the mean individual absolute fitness over a time-step is shorter than a generation, it is the population growth rate (Fisher 1930). The strength of selection is consequently influenced by the population growth rate, as long as the covariance between phenotypic trait values and individual absolute fitness does not vary systematically with the population growth rate.

A dynamical consequence of selection being partially determined by the covariance between phenotypic trait values and individual absolute fitness, is that if the mean of one quantity changes, the mean of the other will also change. This is true as long as i) the function linking the two quantities does not change with time, and ii) there is variation in each quantity within the population. For example, evolution of a phenotypic trait will cause an increase in mean absolute fitness (i.e. the population growth rate) (Fisher 1930). Under these scenarios, both eco-evolutionary dynamics and feedbacks will occur (Coulson et al. 2017). This is because i) the mean of the phenotypic trait and the population growth rate will evolve together, ii) an increase in the population growth rate will increase the size of the denominator in the selection differential, iii) this will reduce the strength of selection which will, in turn, iv) reduce the rate of evolution of the population growth rate. The speed of the eco-evolutionary dynamics, and the magnitude of the eco-evolutionary feedback, both depend upon the association between phenotypic trait values and individual absolute fitness, the phenotypic variance, and the form of the map between genotype and phenotype.

In artificial settings it is possible to control i) the association between phenotypic trait values and individual absolute fitness, ii) the phenotypic variance, and iii), to some extent, the way that environmental variation might influence the genotype-phenotype map (Falconer 1960). Such control does not exist in the natural world, where density-dependent feedbacks and environmental variation can generate temporal change in each of these three quantities. For example, if evolution occurs while the association between phenotypic trait values and individual absolute fitness remains constant in time, populations grow hyper-exponentially (Witting 2002). This is not observed in nature because density-dependent processes such as food limitation and predation prevent long-term hyper-exponential (or even exponential) population growth. Negative density-dependence operates to lower the elevation of the association between phenotypic trait values and individual absolute fitness, which in turns reduces mean individual absolute fitness (the population growth rate) (Coulson et al. 2017).

We use evolutionarily-explicit, density-dependent, structured population models to examine how environmental variation can alter the form of the association between phenotypic trait values and individual absolute fitness, the genotype-phenotype map, and the phenotypic variance, thereby impacting eco-evolution and potentially leading to eco-evolutionary feedbacks.

## THEORETICAL UNDERPINNINGS

Understanding eco-evolutionary dynamics and feedbacks requires dynamic models. However, standard evolutionary theory widely used by empiricists is static, in that it only provides predictions over a single time-step (Falconer 1960, Morrissey et al. 2010). Nonetheless, this static theory is useful as selection differentials on which it is based provide exact descriptions of the strength of observed selection on phenotypic traits (Lande and Arnold 1983), and the Price equations builds on this to provide a complete description of evolution (Price 1970). If models of eco-evolution are to link with the understanding these static models have provided, they must predict time series of components of these static models (Coulson et al. 2010, Coulson et al. 2017, Coulson et al. 2018). We consequently start by considering these models.

We start with the Breeder’s equation. Following standard quantitative genetic theory, we assume that an individual’s phenotypic trait value *z_t_* can be decomposed into a breeding value *A_t_* and an environmental component *E_t_*, so that *z_t_* = *A_t_* + *E_t_*. Next, we write the Breeder’s equation in terms of individual absolute fitness *w_t_* (Heywood 2005) and assume an annual life history and an annual population census such that viability selection can be ignored and mean absolute fitness is equal to the mean recruitment rate 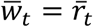, where 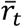 is defined as the per capita production of offspring between times *t* and *t* + 1 that survive to enter the population at time *t* + 1

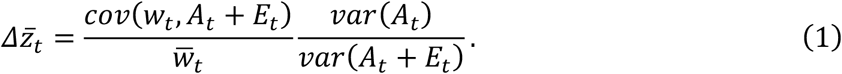

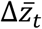 is the change in the mean phenotypic trait value over a generation (=time-step for our annual life history) (Lush 1943), the first term on the right-hand side is a phenotypic selection differential, and the second is the narrow-sense heritability *h*^2^. The variance in the breeding values at time *t* is denoted *var*(*A_t_*) – the additive genetic variance. Equation *(1)* assumes that change in *Ā_t_* caused by selection within the parental generation is transmitted to the offspring generation, and that change in *Ē_t_* is not.

We can rewrite equation (*1*) as 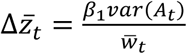 where *β*_1_ is the slope of a linear regression between individual absolute fitness and individual phenotypic trait values (the derivative 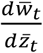). Note that this makes *β*_1_ different from a selection gradient (Lande and Arnold 1983), which is computed using relative fitness values (e.g. 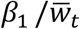 in our notation).

In this formulation, 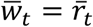 is mean fitness and the population growth rate. Assuming that *β*_1_ and *var*(*A_t_*) do not systematically vary with 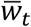, it is apparent that the rate of evolution 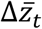 and the population growth rate 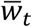 are intimately linked. For example, if we assume a linear fitness function *w_t_* = *β*_0_ + *β*_1_*z_t_* we can write:

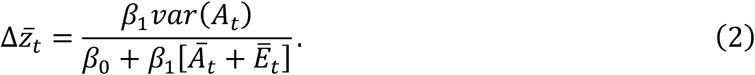

The equations above assume that selection is the only process that alters *Ā_t_* or *Ē_t_* from one generation to the next. This is one reason why the Breeder’s equation does not provide a complete description of evolution of the mean of a phenotypic trait. Non-genetic inheritance and phenotypic plasticity, that can influence the value of *Ē_t_* and contribute to evolutionary dynamics, are processes that are not included in the Breeder’s equation. Put another way, the Breeder’s equation assumes that the genotype-phenotype map does not vary with time due to processes that can alter *Ē_t_*. These processes can be captured, along with selection, with the Price equation (Price 1970).

The Price equation provides a complete description of the change in the mean of a phenotypic trait over a generation. Continuing to assume *z_t_ = A_t_ + E_t_*,

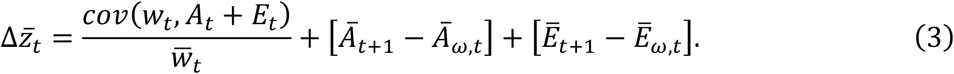

*Ā_ω,t_* and *Ē_ω,t_* respectively describe the mean of the breeding value and environmental component of the phenotype among selected parents.

The first term on the right-hand side is a selection differential. The second term describes how imperfect genetic inheritance (e.g., mutation, recombination) could alter the mean breeding value between offspring and their parents, and the third how non-genetic inheritance (e.g., epigenetics) alters the mean environmental component of the phenotypic trait between offspring and their parents.

The Breeder’s equation is a special form of the Price equation that assumes that [*Ā*_*t*+1_ — - *Ā*_*ω,t*_] = 0 and 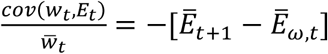. Biologically this means that genetic inheritance does not change *Ā_t_* (i.e. there is no transmission bias), and that *Ē_t_* at birth remains identical in each generation. The first of these assumptions is reasonable for large populations; the second is justified if the environmental component of the phenotypic trait is determined by completely random developmental noise (Falconer 1960) and not by any variation in the environment. If the environmental component of the phenotypic trait is dependent upon non-random aspects of the external environment (including non-genetic inheritance), then the assumption will be violated (Kruuk et al. 2002). If this is the case, the Price equation is required to describe evolution.

In an annual life history 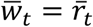; in a non-annual life history, the population growth rate and mean fitness over a time-step are the sum of mean survival and mean recruitment, 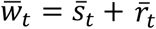. Replacing 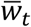 with 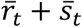 and expanding *(3)* we can write the Price equation in terms of fitness and transition functions acting through survival, recruitment, development and genetic and non-genetic inheritance on the distribution of z. In this form, the Price equation applies to iteroparous species:

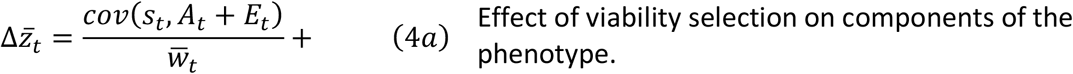

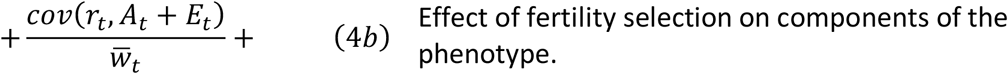

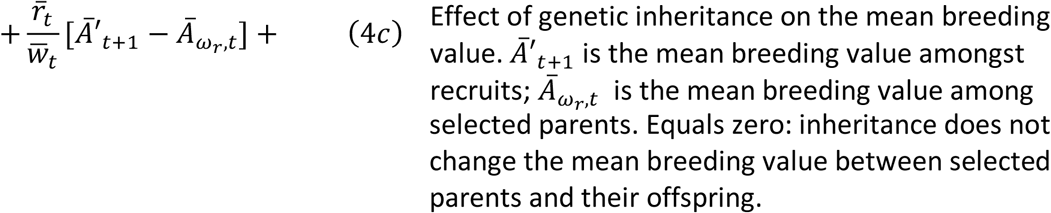

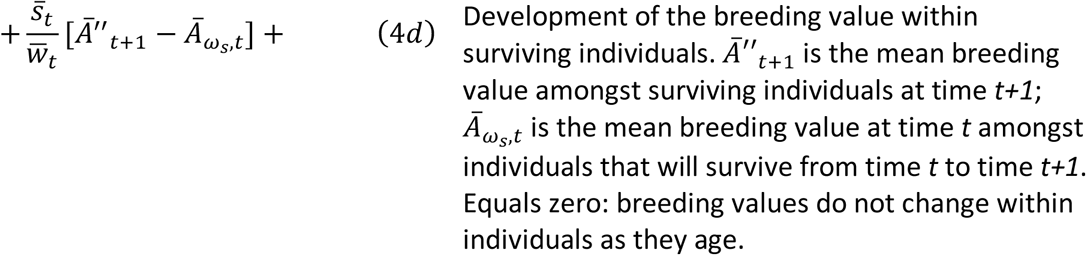

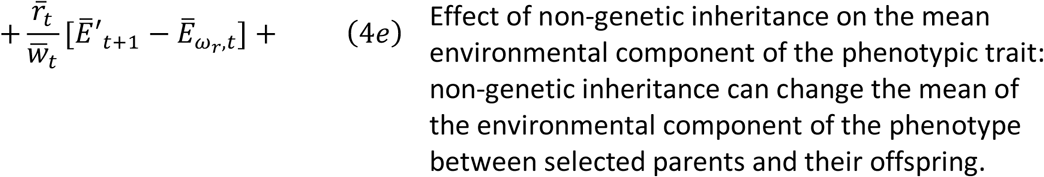

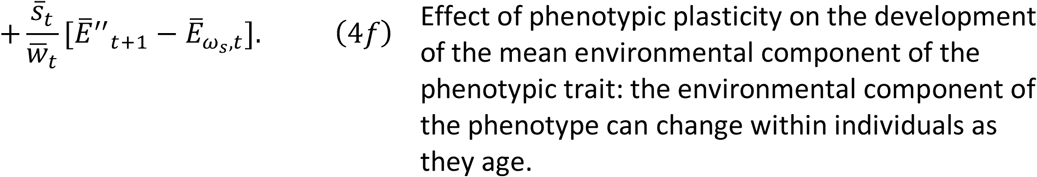

This provides a complete description of all of the processes that can influence change in the mean phenotype in an iteroparous species over a time-step. The formulation does assume *z_t_ = A_t_ + E_t_* and that all processes remain constant with age. The latter assumption could be relaxed using the approach in (Coulson and Tuljapurkar 2008). Note that some authors have further decomposed *cov*(*r_t_,A_t_* + *E_t_*) into *cov*(*r_t_,A_t_*) + *cov*(*r_t_, E_t_*) and assumed that these two quantities can change independently of one another (Bonnet et al. 2017). We do not do this here as this violates the additivity assumption of *z_t_ = A_t_ + E_t_* of quantitative genetics (Appendix 1) that we are currently employing.

Each of the terms in the static Breeder’s and Price equations are emergent properties of the distribution of the phenotypic trait (and its components) at time *t,* fitness functions determining terms in equation *(4a)* and *(4b)*, and transition functions determining terms in equations *(4c)* to *(4f)* (Coulson et al. 2010, Childs et al. 2016). Integral projection models (IPMs), which calculate changes in the distribution of a continuous trait by integrating over a series of fitness and transition functions (Coulson et al. 2010, Childs et al. 2016) provide a framework with which to make the Price and Breeder’s equations dynamic. To do this, it is consequently necessary to specify these functions (Coulson et al. 2017) that can then be combined into a model to iterate forward the dynamics of the distribution of *Z_t_* = *A_t_* + *E_t_*.

In the case of an annual life history, a reproductive fitness function, *R*(*A_t_* + *E^t^*), is used to map the bivariate distribution of breeding values and environmental components of the phenotype at time *t, n*(*A_t_, E_t_*), to a new distribution at time *t* + 1, 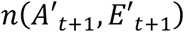 (Ellner et al. 2016),

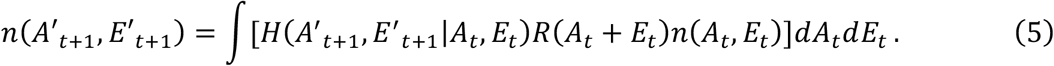

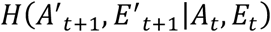 is a transition function that describes the probability of transitioning from trait value *A_t_* + *E_t_* at time *t* to all possible trait values 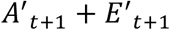 at time *t* + 1 (Easterling et al. 2000). Trait value *Z_t_ = A_t_ + E_t_* is expressed in the focal individual, while trait values 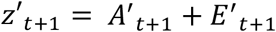 are expressed in offspring. The transition function is needed when predicting evolutionary change over multiple time-steps if offspring that recruit to the population at time *t* + *1* and their parents at time *t* do not perfectly resemble one another.

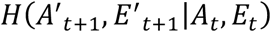 can be formulated to capture the second term on the right-hand side of the Breeder’s equation *(1)* or the last two terms on the right-hand side of the Price equation (*3*), and so can describe genetic and non-genetic inheritance. A dynamic form of the Breeder’s equation (*1*) requires 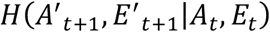 to be specified such that it produces a distribution 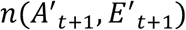 that has means of 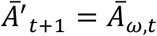 and 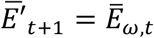 and a variance-covariance matrix 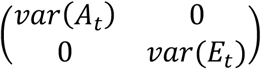 where *var*(*A_t_*) can be selected to be a constant, or to change with time as a function of selection, and *var*(*E_t_*) is constant (Childs et al. 2016, Coulson et al. 2017, Rees and Ellner 2019). There are a number of ways this can be achieved (Appendix 2). Any formulation of 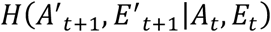 that results in 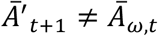 and/or 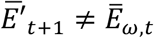 provides a dynamic form of the Price equation *(3, 4)*. Once the transition and fitness functions are parameterised, derivations from Coulson *et al.* (2010) can then be used to create time series of each of the components in the various forms of the Price and Breeder’s equations described above.

Equation (*5*) can be extended to incorporate per-time-step survival and recruitment functions to allow models of iteroparous species. Models now include two transition functions: a development function 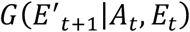 operating on surviving individuals, and an inheritance function 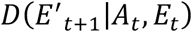 that captures the similarity between parental and offspring traits (Easterling et al. 2000, Ellner and Rees 2006, Coulson 2012). The development function operates with a survival fitness function *S*(*A_t_* + *E_t_*) that determines the strength of viability selection, while the inheritance function operates with the recruitment fitness function *R*(*A_t_ + E_t_*) that determines the strength of fertility selection:

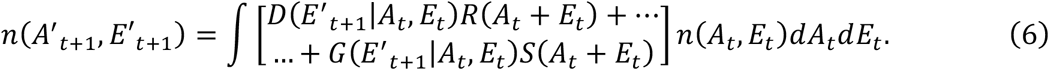

Each process that can contribute to the change in the mean phenotype in an iteroparous species (i.e. Equation(4)) is captured in this IPM: term (*4a*) is determined by *S*(*A_t_* + *E_t_*)*n*(*A_t_, E_t_*), term (*4b*) by *R*(*A_t_* + *E_t_*)*n*(*A_t_, E_t_*), terms (*4c*) and (*4e*) by 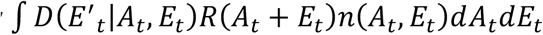 and terms *(4d)* and *(4f)* by 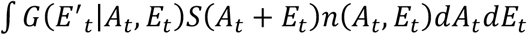 (Childs et al. 2016).

## METHODS

Our aim here is to incorporate environmental trends and feedbacks caused by densitydependence into the functions in equations (*5*) and (*6*). We will examine how this impacts the association between phenotypic trait values and individual absolute fitness, the phenotypic variance, and the map between genotype and phenotype, and how, ultimately, this generates temporal dynamics in the components of the Breeder’s and Price equations, rates of evolution, phenotypic trait dynamics, and population growth.

We start with a model of the form of equation (*5*) with the phenotypic trait separated into its additive genetic and environmental components, *z_t_ = A_t_ + E_t_*, but where fitness is not separated into its survival and recruitment components. We assume a linear association between phenotypic trait values and individual absolute fitness and iterate *n*(*A_t_, E_t_*) given the following parameterisation,

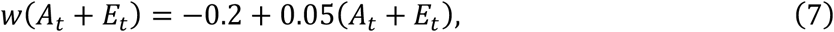

and a starting bivariate normal distribution *n*(*A*_*t*=1_, *E*_*t*=1_) with means *Ā_t_ = Ē_t_* = 10 and a variance-covariance matrix 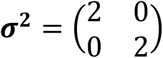. The choice of parameter values is for illustration rather than representing any particular system. The mean of the breeding value distribution does not change between parents and their offspring.

We ran four model scenarios. In scenarios 1) and 2) the mean and variance of the environmental component of the phenotype remained constant across generations, while in scenarios 3) and 4) the mean of the environmental component declined with time such that *Ē_t_* = 10 ~ 0.05*t*. In scenarios 1) and 3) selection eroded additive genetic variation, while in scenarios 2) and 4) it did not. Scenarios 1) and 2) are dynamic forms of the Breeder’s equation, while scenarios 3) and 4) are dynamic forms of the Price equation. The last two scenarios capture a process where some aspect of the environment can determine the dynamics of the environmental component of the phenotype (Kruuk et al. 2002).

The linear assumption of the simple model is unrealistic because litter size and survival rates are bounded. In addition, in variable environments, survival, recruitment, development and inheritance functions for the environmental component of the phenotype depend on environmental factors such as the population size (Kruuk et al. 2002, Bassar et al. 2017). Moreover, although breeding values remain constant with age, the environmental component of the phenotypic trait can change within individuals - phenotypic plasticity (Falconer 1960). We consequently developed a more complex, density-dependent model, where fitness is separated into its survival and recruitment components, and where the environmental component of the phenotype can develop within surviving individuals, and can vary between parents and their offspring. We took a published density-dependent Integral Projection Model (IPM) of the dynamics of body size from females in an individually marked population of Soay sheep *(Ovis aries)* living on the island of Hirta in the St. Kilda archipelago in Scotland (Coulson 2012). The published IPM was modified to be of the form of equation *(6)*. Functions and parameter values are provided in Appendix 3.

We selected the starting population structure of each evolutionarily explicit simulation by using the equilibrium population structure from an ecological simulation where the additive genetic variance was set to 0 so evolution cannot occur. Each simulation was run for 100 time-steps.

As well as the baseline model, in which the survival, recruitment, development and inheritance functions were all density-dependent, we ran four simulations where we set the densitydependence parameter to 0 in either the survival fitness function, the recruitment fitness function, the development function, or the inheritance function for *E_t_*. Although this is a large perturbation, it reveals interesting dynamics whereby perturbing one function influences the role of other functions on eco-evolution.

## RESULTS

In our evolutionarily explicit IPM with a linear fitness function, evolution, defined as a change in the mean breeding value Δ*Ā_t_*, was fastest when additive genetic variance was not eroded by selection, and when, simultaneously, there was a negative temporal trend in the mean environmental component of the phenotype *Ē_t_* (Figure 1(A)). The negative temporal trend in *Ē_t_* accelerated the rate of evolution by reducing the population growth rate 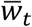 (Figure 1(B)). In spite of accelerating the rate of evolution, the trending values of *Ē_t_* with time slowed the rate of phenotypic change, as positive change in *Ā_t_* was countered by negative change in *Ē_t_* (Figure 1(C)). The population growth rate 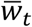, which is also mean fitness, increased fastest when additive genetic variance was not eroded, and *Ē_t_* did not decline with time (Figure 1(C)). As expected, linear selection reduced the additive genetic variance if it was not replaced in each generation (Figure 1(D)). When there was no temporal trend in *Ē_t_*, the association between the mean breeding value and the population growth rate was linear (Figure 1(E)), while it was slightly curvilinear when *Ē_t_* declined with time. As the association between phenotypic trait values and individual absolute fitness was linear, there was, by definition, a linear association between the mean phenotypic trait and the population growth rate, with a slope of *β*_1_ (Figure 1(F)).

**Figure 1.**
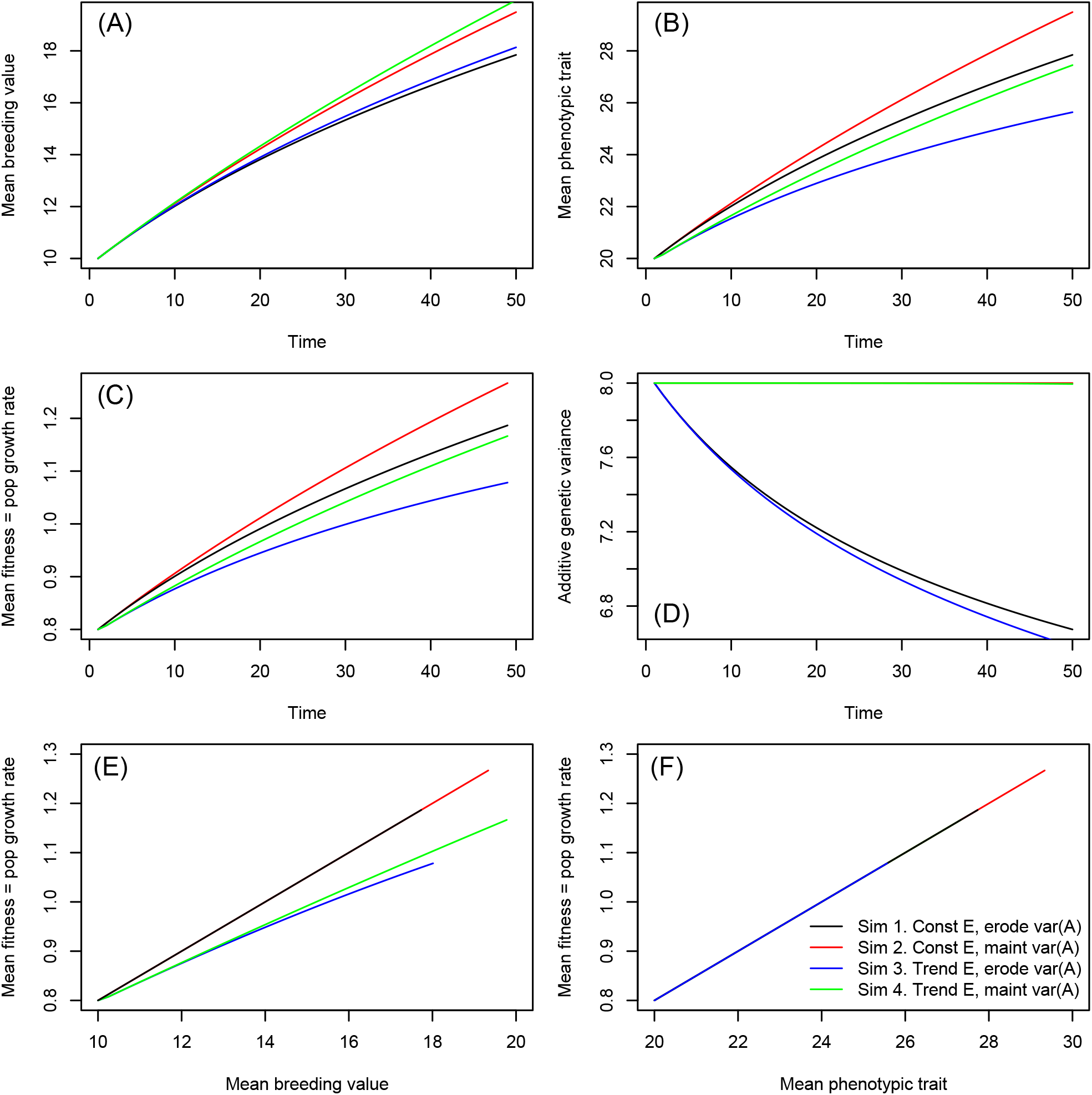
Evolution of (A) the mean breeding value, (B) the mean phenotypic trait, (C) mean fitness, and (D) the additive genetic variance in simple models over 50 generations. The black line represents sim 1), where the additive genetic variance is eroded and the mean and variance of the environmental component of the phenotypic trait remain constant with time. The red lines represent sim 2), where additive genetic variance is not eroded by selection, and the mean and variance of the environmental component of the phenotypic trait remain constant with time. The blue (sim 3)) and green line (sim 4)) are the same as sim 1) and sim 2), except we impose a weak temporal downward trend on the mean environmental component of the phenotypic trait. (E) In models with no temporal trend in the environmental component of the phenotypic trait, the relationship between the mean breeding value and mean fitness is linear, while it is slightly curvilinear in models with temporal trends in *E*. In contrast, there is (F) a linear association between the mean phenotypic trait and mean fitness in all four models.

We can combine these insights to graphically illustrate the evolutionary, phenotypic, and population-growth dynamics our results reveal with a plot of *A_t_* against *E_t_* that includes fitness clines (Figure 2(A)). Each diagonal dashed line represents equivalent values of the phenotypic trait *Z_t_* made from different combinations of *A_t_* and *E_t_*. Because the association between phenotypic trait values and individual absolute fitness was linear and included no additional terms beyond *β*_0_ and *β*_1_, these dashed lines also represent the population growth rate. The darker the colour of the line, the larger the value of *z_t_* and the population growth rate. The central solid line contours represent the bivariate distribution of *A_t_* and *E_t_* at time *t* = 0 when simulations are begun. The green, red and purple arrows represent phenotypic change. When evolution was cryptic (blue arrow), meaning that neither 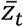, nor 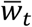, changed, *Ā_t_* increased at the same rate as *Ē_t_* decreased such that Δ*Ā_t_* - Δ*Ē_t_* = 0. Because 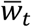 did not change in this situation, evolution proceeded rapidly as the denominator in the selection differential remained constant with time. Dynamic forms of the Price equation are required to capture such cryptic evolution. The green line represents dynamics predicted by dynamic forms of the Breeder’s equation, with *Ā_t_* and 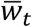 evolving, thus resulting in phenotypic change, but *Ē_t_* remaining constant with time. Evolution occurred more slowly than when it was cryptic because 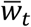 increased with time, increasing the denominator of the selection differential. The red line captures evolutionary dynamics where the mean breeding value and the mean environmental component of the phenotype change in the same direction, increasing both 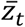 and 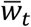. A dynamic form of the Price equation is required to capture such dynamics, and evolution proceeded even more slowly than in the previous two examples, because 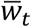 increased even faster. Finally, the purple arrow represents no evolution, but phenotypic change due solely to change in *Ē_t_* – i.e. via non-genetic inheritance (or phenotypic plasticity in iteroparous species). A dynamic form of the Price equation is required to capture such dynamics.

We now turn to the more complex Soay sheep model that extends insights from the linear model in three ways: i) it is for an iteroparous life history, ii) it incorporates density-dependence in the survival and recruitment functions (thus reducing the intercepts of the fitness functions with time as evolution proceeds), and iii) it has non-linear fitness functions (meaning that the slope of the association between phenotypic trait values and individual absolute fitness will vary as evolution proceeds).

**Figure 2.**
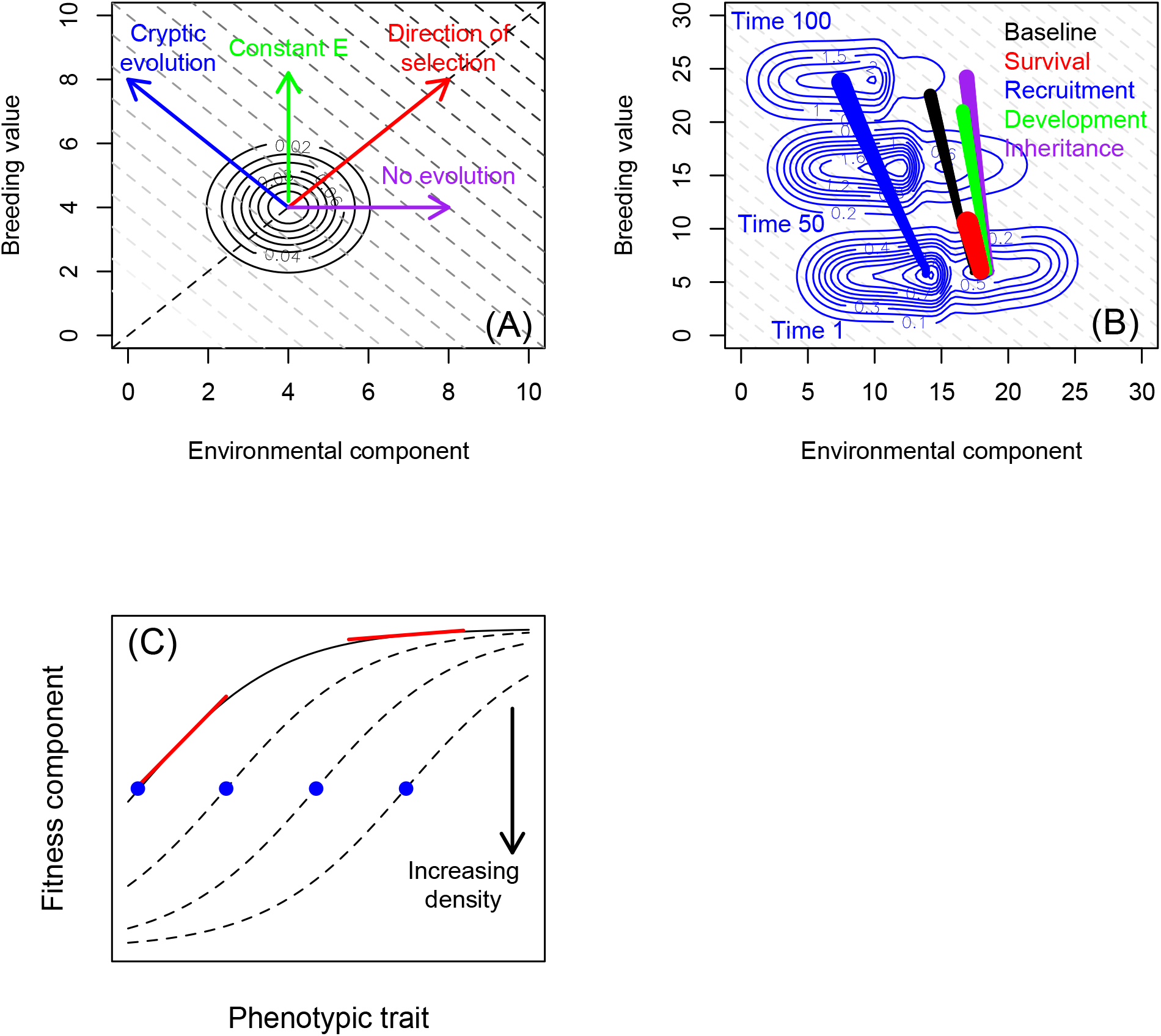
The dynamics of phenotypic traits and mean fitness. (A) Because the breeding value and the value of the environmental component of a phenotypic trait sum to produce the phenotypic trait, there are multiple ways the same phenotypic trait value can be produced (dashed grey lines). If the phenotypic trait linearly influences fitness, these phenotypic trait clines also represent fitness clines, generating linear fitness contours. Larger values of the trait, and of mean fitness, are depicted with darker dashed lines. Depending upon the dynamics of the environmental component of the phenotype, selection can fail to climb across contours, or can climb at a range of rates even for the same, constant, derivative of mean fitness to the mean phenotype (i.e. phenotypic slope in the linear fitness function). Because mean fitness remains lowest when evolution is cryptic, all other things being equal, selection will also be strongest, and evolution fastest. Cryptic evolution occurs when non-genetic inheritance or phenotypic plasticity impact the environmental component of the phenotype in a manner that exactly counters gains to the additive genetic component of the phenotype attributable to selection. (B) The bivariate dynamics of the mean breeding value and the mean environmental component of the phenotypic trait over time in each simulation for the modified sheep model. The starting point of each simulation is towards the bottom of the figure. The increasing width of the thick lines represent how population size (as a consequence of the dynamics of mean fitness) changes as the simulation progresses. Bivariate distributions of the components of the phenotypic trait are drawn as contour plots at time *t*=1, *t*=50 and *t*=100 for the simulation where density-dependence was removed from the recruitment (fitness) function. Evolution is captured by the length of the thick line in the y-axis, while change in the environmental component of the phenotypic trait is captured by change in the x-axis. (C) The derivative of mean fitness to the mean phenotype (red line) can vary as either the mean phenotypic trait increases when fitness functions are non-linear (blue dots), or when environmental drivers such as increasing population density lower the intercept of the fitness function (dashed black lines).

Figure (2(B)) shows the joint dynamics of *Ā_t_* and *Ē_t_* for each of the simulations of body size in female Soay sheep. We do not include fitness clines such as those in Figure (2(A)) because the incorporation of density-dependence complicates the association between 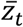 and 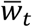. Before we explore these results in further detail, it is helpful to appreciate that change in 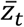 can alter both the elevation (blue dots) and slope (red lines) of the expected association between phenotypic trait values and individual absolute fitness (red lines; Figure (2(C))). The way a system evolves depends on both the rate that phenotypic change alters the location of the distribution of phenotypic trait values on a fitness function, and the rate at which evolution of the population growth rate alters the form of the fitness function itself.

Compared to the baseline model where all four functions were density-dependent, removing density-dependence from the recruitment and inheritance functions in the modified sheep model increased the rate of evolution of body size, while removing it from the survival and development function decreased it (Figure 2(B)). By far the largest effect was caused by perturbing the survival function.

Temporal changes in *Ē_t_* were negative in all models, with the greatest change observed when density-dependence was removed from the recruitment function (Figure 2(B)). This negative trend only partially countered the positive trends in *Ā_t_* (Figure 3(A, B)), such that 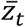 increased with time in all models (Figure 3(B)), but to a lesser extent than *Ā_t_* (Figure 3(A)).

**Figure 3.**
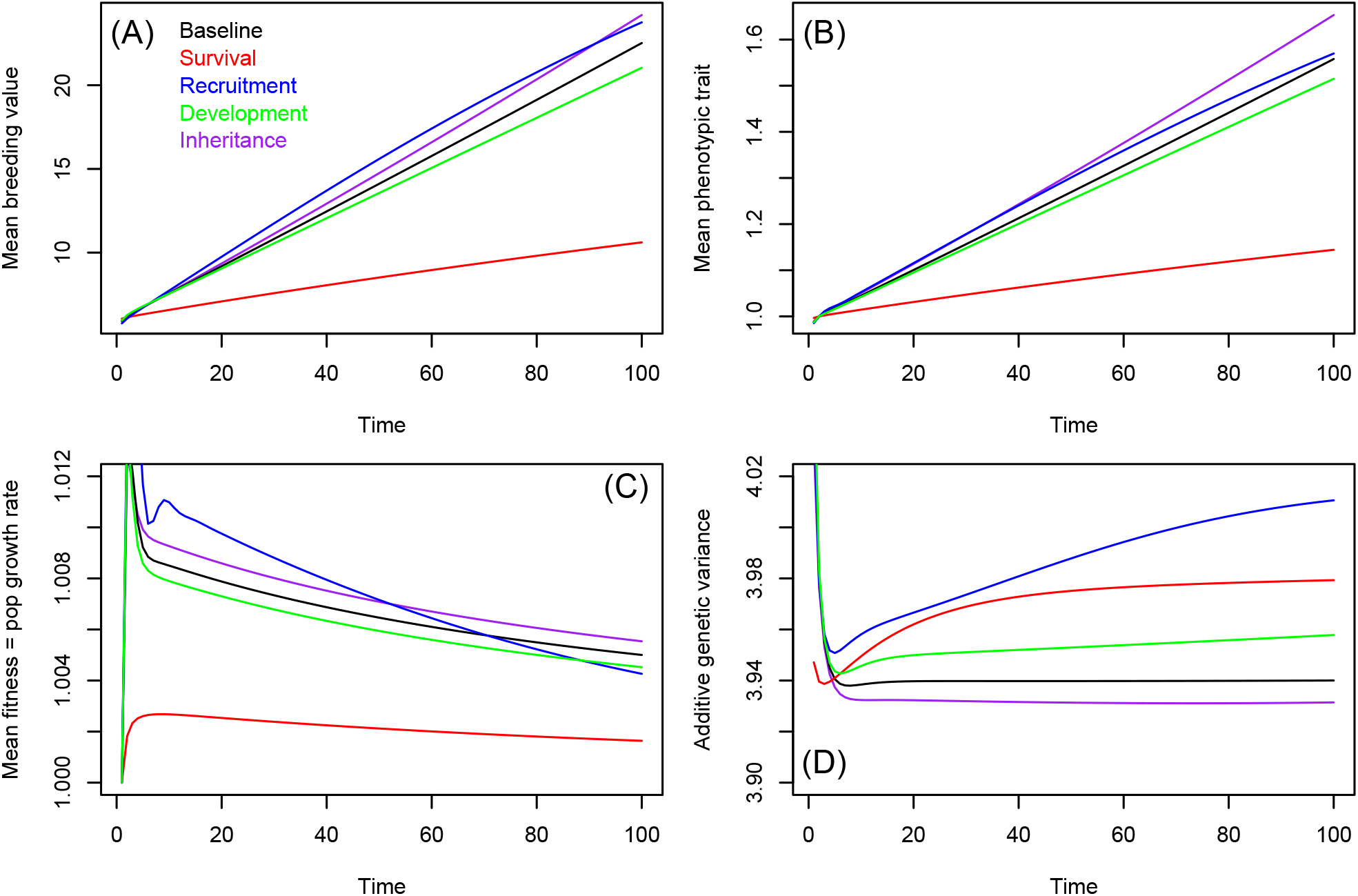
Evolution of (A) the mean breeding value, (B) the mean phenotypic trait value, (C) mean fitness, and (D) the additive genetic variance in the modified sheep model. The baseline model (black lines) has negative density-dependence in each function, while the coloured lines represent simulations where density-dependence has been removed from one particular function.

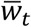 evolved at similar rates in the baseline model and models where density-dependence was removed from the recruitment, inheritance and development functions (Figure 3(C)). In contrast, 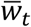 evolved much more slowly when density-dependence was removed from the survival function (Figure 3(C)), although initial population size was much larger in the density-independent survival model than in any of the other models (Figure 2(B)). The additive genetic variance increased slightly with time when density-dependence was removed from the survival, recruitment and development functions, but remained constant when inheritance was density-independent, as well as in the baseline model (Figure 3(D)).

Removing density-dependence from each fitness function impacted evolutionary, phenotypic and population-size change via a number of different routes. First, removing density-dependence resulted in different starting population sizes and structures (Figures 4(A)-(E)). In the baseline model and the model with density-dependence removed from the development function, there was clear bimodality in the distribution of *Z_t_* throughout the simulation (Figures 4(A, E)). The bimodality reflects a population with similar proportions of smaller juveniles and larger adults. There was weaker evidence of a bimodal distribution at time *t* = 1 for the other three models, with bimodality weakening as the simulations proceeded (Figures 4(B-D)). In models with densityindependent survival and inheritance, small juveniles were comparatively rare from the start, and became rarer still over time. In contrast, in the density-independent recruitment model, the population was initially dominated by small juveniles, but then also became increasingly dominated by larger adults as the simulation proceeded.

**Figure 4.**
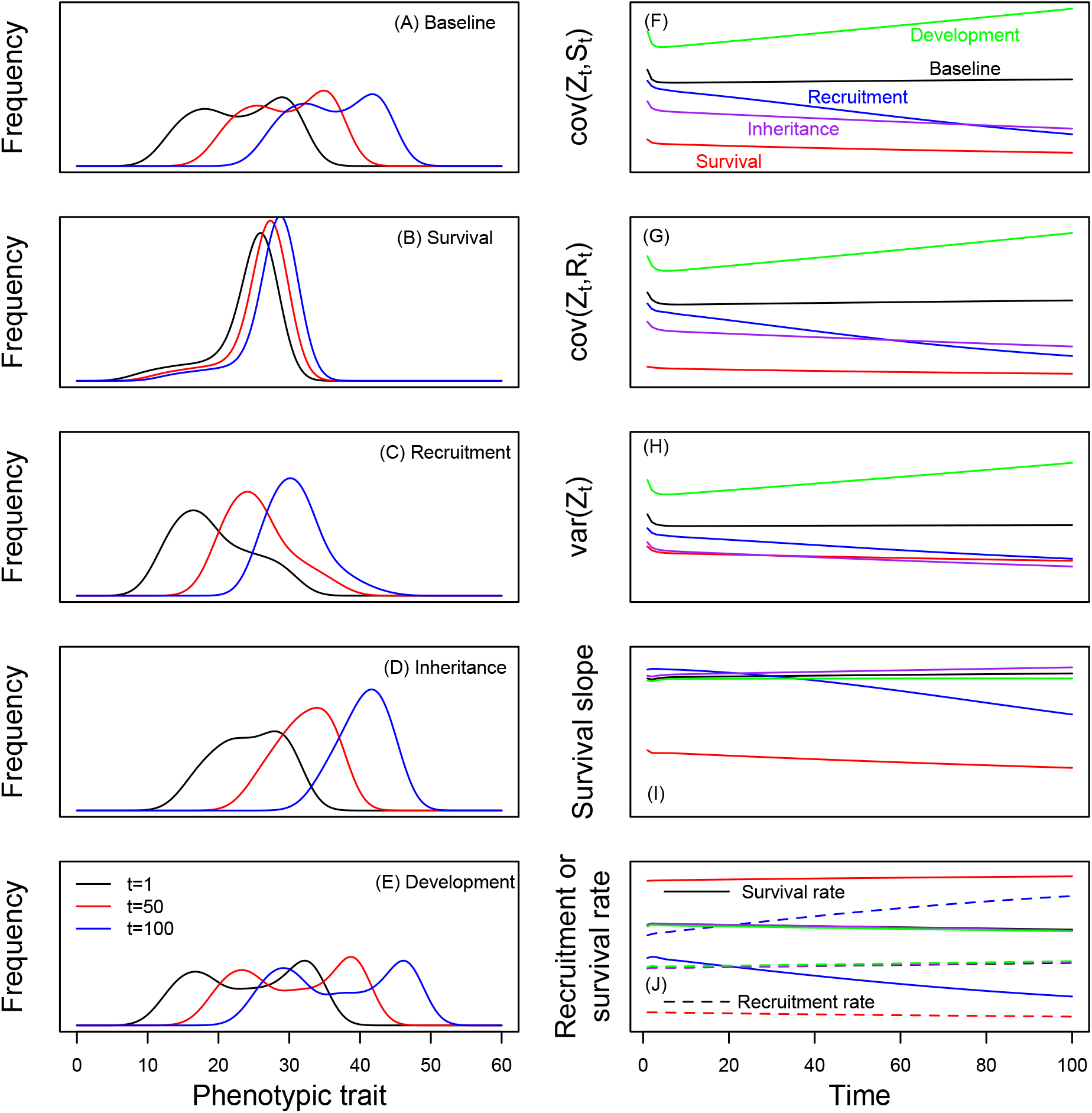
Dynamics of the phenotypic trait distribution in the baseline model (A) and in models with negative density-dependence removed from each function (B-E). Distributions are drawn at times *t*=1, *t*=50 and *t*=100. These dynamics are determined in part by selection (the derivative of mean fitness to the mean phenotypic trait value) that consists of (F, G) covariances between the trait and components of fitness, and (H) the phenotypic variance in the trait. (I) derivative of mean survival to the mean phenotype (similar patterns for recruitment not shown) and (J) the temporal dynamics of mean survival and recruitment rates.

The rates of evolution of the phenotypic trait and mean fitness are determined by i) *β*_1_, ii) 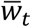 (which depends upon *n*(*z_t_*) and the fitness functions), and iii) *var*(*A_t_*). Because the models were density-dependent, the mean population growth rate tended towards unity, and differed little between simulations (Figure 3(C)). Relatively little of the dynamical difference between the models was consequently due to differences in 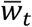 across simulations. Similarly, because the additive genetic variance remained approximately constant in all simulations (Figure 3(D)), this too was not a major driver of the dynamics we report from the sheep model.

Most of the dynamical differences between the simulations were consequently caused by processes generating variation in *β*_1_: the covariance between individual phenotypic trait values and individual absolute fitness and the phenotypic variance. These two terms tended to show similar temporal dynamics (Figure 4(F-H)). Consequently, the derivative between mean survival and the phenotype (the strength and direction of viability selection) remained approximately constant in the baseline model and the density-independent development and inheritance models, while it trended downwards as the simulation proceeded in the density-independent survival and recruitment models (Figure 4(I)). Similar patterns were observed for the derivative of mean recruitment to the mean phenotype (not shown).

These temporal changes in the derivatives of the mean fitness components to the phenotypic trait value resulted in approximately constant survival and recruitment rates in the density-independent survival model, weak decreases in survival rates and weak increases in recruitment rates in the baseline, density-independent development and density-independent inheritance models, and stronger temporal trends for decreased survival and increased recruitment rates in the density-independent recruitment model (Figure 4(J)). In other words, the patterns we report in mean fitness were caused by opposing trends in fitness components in these simulations.

The dynamics of *β*_1_ that generated these contrasting patterns across the different simulations were determined by the shape of the fitness function (Figure 2(C)). As density increased, the intercept of the logistic fitness functions became smaller due to the density-dependence terms in the models. In contrast, as mean body size increased, the body size distribution moved to the right on the x-axis of figure 2(C). To some extent, these two processes tended to cancel one another out, such that mean survival and mean recruitment tended to change relatively slowly. The one exception to this pattern is in the density-independent recruitment model, where increasing body size increased recruitment rates but did not lower the intercept of the function. Instead, mean survival rates decreased as density increased because mean fitness evolved.

To sum up, our results show that the Soay sheep population is primarily regulated via density-dependent survival. When density was removed from the survival function, the population consequently increased to a large size, in turn reducing recruitment rates, development rates, and the mean value of the environmental component of the phenotype. Large individuals, however, had very high survival, at a point when the derivative of mean survival to body weight was very shallow – i.e. *β*_1_ was close to zero and evolution of both the mean phenotypic trait and mean fitness was slow.

## DISCUSSION

The structured, evolutionarily-explicit models we developed to examine the dynamics of terms in the Breeder’s and Price equations build on existing approaches by Lande (Lande 1982), Barfield et al. (Barfield et al. 2011), Childs et al.(Childs et al. 2016), Coulson et al. (Coulson et al. 2011, Coulson et al. 2017) and Rees and Ellner (Rees and Ellner 2019). The approach is powerful as it allows the simultaneous study of population dynamics and the evolution of phenotypic traits, life histories, and mean fitness (Coulson et al. 2011). In this paper we use dynamic models to generate dynamics of components of the Breeder’s and Price equations.

Population dynamics and phenotypic evolution are both determined by three drivers: i) the distribution of phenotypic trait values, ii) the fitness functions that determine how expected survival and reproductive rates vary with phenotypic trait values and the environment, and iii) details of the genotype-phenotype map (Lande 1982, Lynch and Walsh 1998). The latter can include phenotypic plasticity, and genetic and non-genetic inheritance (in all its forms). Our models reveal how rates of evolution (defined as Δ*Ā_t_*), of phenotypic change, and of change in the population growth rate, are determined by factors that influence each of these three drivers.

In the absence of any processes that alter these drivers with time, genetic change, phenotypic change, and population growth rate change will occur simultaneously (Witting 2002). In the case of a linear association, the rate of change of each will slow with time. This is an example of eco-evolutionary feedbacks. If the association between the phenotypic trait and fitness is exponential, then the rate of change is constant. This is because the slope of the association between the phenotypic trait and fitness changes with time as the mean phenotype evolves.

In natural systems, because populations cannot increase in size indefinitely, the association between the phenotypic trait and fitness must be influenced by processes such as densitydependence or environmental change, to prevent the population growing hyper-exponentially. These processes are typically modelled as being additive (Ellner et al. 2016), and act to reduce the intercept of the association between the phenotypic trait and fitness, as evolution increases the mean value of the phenotypic trait (Smallegange and Coulson 2013). However, these processes can also interact with the phenotypic trait to alter the slope of the association. In our models, the survival and fertility (fitness) functions can capture all these processes (Coulson et al. 2017).

Environmental drivers such as density-dependence and climatic variation can influence the rate of evolution, phenotypic change, and the population growth rate, by directly impacting the phenotypic trait via processes other than selection - e.g. phenotypic plasticity and non-genetic inheritance (Nussey et al. 2007, Reed et al. 2011). These processes can affect the rate of evolution, phenotypic trait dynamics, and the ecology of a system. In particular, these drivers can alter the mean or variance of the trait, by changing the environmental component of the trait (Coulson et al. 2017). In our models, these effects are captured by the development and inheritance functions - where the latter also captures genetic inheritance.

In natural systems, environmental drivers typically impact fitness, development and inheritance functions simultaneously, and potentially in different ways in different ages and sexes (Simmonds et al. 2019). A consequence of this is that the environment can simultaneously influence evolution, phenotypic trait dynamics, and the population growth rate (e.g. ecology) in multiple, interacting ways (Coulson et al. 2011). For example, density-dependence can alter the association between a phenotypic trait and fitness, and alter the mean and variance of the environmental component of the phenotype.

The Breeder’s equation was developed as a model to predict evolution over a single time-step in artificial environments when humans specify the association between the phenotypic trait and fitness to produce a desired number of offspring from those individuals they allow to reproduce (Lush 1943, Falconer 1960, Lynch and Walsh 1998). In addition, they minimise the effect of environmental drivers on the phenotypic trait by typically providing *ad libitum* food and removing predation- and disease-related causes of death. Under these circumstances, the Breeder’s equation performs well. Yet, when it has been used as a retrospective tool to explain evolution in natural systems it has typically failed (Merilä et al. 2001). This is because processes that can alter the rate of evolution in natural settings are not included in the model, and are treated as nuisance variables to be corrected for in statistical analyses (Kruuk 2004). However, these processes can influence evolution (and ecology) and should consequently be included in predictive models if they are found to explain significant variation in statistical analyses used to estimate additive genetic variances and the strength of selection. Our approach allows for these processes to be explicitly incorporated into predictive models (Childs et al. 2016, Coulson et al. 2017, Simmonds et al. 2019).

The Breeder’s equation does not incorporate all the processes that can generate phenotypic change (Morrissey et al. 2010). This observation limits the utility of estimates of additive genetic variances and heritabilities from free-living populations (Kruuk 2004), because, if these processes are important, they should be incorporated into predictive models. The Price equation, on the other hand, does provide an exact description of phenotypic evolution (Price 1970). However, it is a tautology (Heywood 2005), and simply describes the difference between two means. It is consequently a very useful tool to retrospectively decompose phenotypic change into all factors that contributed to it (Coulson and Tuljapurkar 2008), but it is not dynamically sufficient, and consequently cannot be used to make predictions of future dynamics. The structured population models we develop are dynamically sufficient, and provide a dynamic form of the Price equation (Childs et al. 2004, Coulson et al. 2010). In previous work, we have demonstrated how these models can be parameterised for real systems (Simmonds et al. 2019).

The class of structured models we have developed to predict evolutionary change in natural systems is widely used in ecology (Caswell 2001). A consequence of this is that our models can be used to predict simultaneous ecological and evolutionary change (Coulson et al. 2011). Ecoevolution is frequently described as ecological change generating evolutionary change, which in turn generates ecological change, and so on *ad infinitum* (Hendry 2016). The logic behind this interpretation is that ecology and evolution are two separate processes that can influence one another (Pelletier et al. 2009). Our work challenges this interpretation, as we show that ecological change and evolution occur simultaneously, as a consequence of ecological change or evolution altering fitness, development and inheritance functions (see also (Coulson et al. 2011, Childs et al. 2016, Coulson et al. 2017)).

Our approach is based on the logic that a population can be represented by a multivariate distribution of genotypes and phenotypes (Easterling et al. 2000, Ellner and Rees 2006). The dynamics of this distribution are then determined by survival and reproduction, development and inheritance functions. We consider these four processes as fundamental. The dynamics of the distribution’s volume (size) are the population dynamics, while the dynamics of the mean genotype (and sometimes phenotypic trait) capture evolution (Smallegange and Coulson 2013). These models can also be used to study the dynamics of life history descriptors such as generation length and lifetime reproductive success (Ellner and Rees 2006, Steiner et al. 2012, Steiner et al. 2014), and to examine life history evolution (Childs et al. 2003, 2004). The fundamental processes that these models capture can vary as a function of age, sex, location, and genotype, and they can include aspects of the biotic and abiotic environment such as the size and structure of the focal species or of other species in the community, climatic variation, or variation in resource availability (Ellner and Rees 2006, Coulson 2012, Schindler et al. 2015, Ellner et al. 2016, Bassar et al. 2017). These models consequently offer a flexible framework to explore how populations will ecologically and evolutionarily respond to environmental change.

In this paper we provide two simple models to demonstrate the utility of our approach. However, more complex models can be developed by extending such models. For example, we can relax the assumption of an additive genotype-phenotype map, and extend models for multiple traits, age- and sex-structure (Schindler et al. 2015, Coulson et al. 2017). It is also possible to construct models of interacting species (Bassar et al. 2017). Although there will doubtless be challenges in parameterising some versions of these models, the approach we describe here offers enormous potential to improve predictions of eco-evolution in natural settings.

## ACKNOWLEDGEMENTS

Tom Potter is joint funded by NERC DTP and Lamb and Flag studentships at Oxford University, and Anja Felmy is funded by an Early Postdoc Mobility Fellowship from the Swiss National Science Foundation (P2EZP3_181775). We thank Dylan Childs and Joe Travis for helpful comments on an earlier version of the manuscript.

## Appendix 1

A key premise of the standard quantitative genetic approach is that the genotype-phenotype map can be described as *z = A + E* (Falconer 1960). If a phenotypic trait is measured in a particular unit, say length in millimetres, then *A* and *E* also take these units. The second premise is that selection operates on the entire phenotype via a selection differential. If we consider a linear fitness function *E*(*w*) = *β*_0_ + *β*_1_*z* we can also write *E*(*w*) = *β*_0_ + *β*_1_(*A + E*) = *β*_0_ + *β*_1_*A* + *β*_1_*E*. This equation says the effect of a one-unit change on fitness in the additive genetic and environmental components of the phenotype is identical.

A linear regression slope is defined as the covariance between the response and predictor variable, divided by the variance in the predictor variable. Consequently,

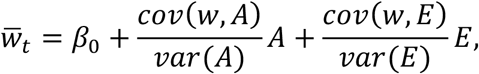

where 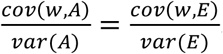. If 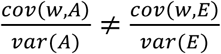 then *z* ≠ *A* + *E*. Instead, the genotype-phenotype map will be of the form *z = A + γE*, where *γ* ≠ 1. In general, this means that any difference in slopes of the breeding values and the environment components of the phenotype with fitness can only occur if the genotype-phenotype map has changed with time during the period over which data were collected to parameterise models.

Some authors have argued that selection can operate in contrasting ways on the environmental and additive genetic components of the phenotype while assuming *z_t_ = A_t_ + E_t_* (Morrissey et al. 2010). In the most extreme cases they argue that selection on the breeding value is in one direction, while selection on the environmental component of the phenotype is in a contrasting direction such that *γ* < 0 while *A_t_* and *E_t_* do not covary (Bonnet et al. 2017). Although mathematically possible, such a genotype-phenotype map seems biologically implausible. We have been unable to identify a biologically plausible genotype-phenotype map where a one-unit increase in *A_t_* will result in an increase in fitness, while a one-unit increase in *E_t_* will result in a decrease, without invoking some dependence of *A_t_* and *E_t_* on one another.

A more plausible genotype-phenotype map would involve phenotypic plasticity, where a breeding value *A_t_* produces a different phenotypic value *z_t_* in different environments. If *z_t_ = A_t_ + E_t_* this requires *E_t_* to vary with the environment *θ*, such that *E* can be made a function of the environment *E*(*θ_t_*). Phenotypic plasticity can then be captured with a number of formulations such as *z_t_ = A_t_* + *E*(*θ_t_*) or *z_t_ = A_t_ + E*(*θ_t_*) + *αA_t_E*(*θ_t_*) (Coulson et al. 2017). In the first formulation, the mean and variance of *E_t_* can vary with the environment, but the genotypephenotype map remains *z_t_ = A_t_ + E_t_* such that the slope between the environmental component of the phenotype and fitness equals the slope between the additive genetic component of the phenotype and fitness, but *Ē_t_* change with time, altering the numerical form of the genotypephenotype map. In contrast, in the second formulation, the breeding value-by-environment interaction means that the slope between fitness and each component of the phenotype differs. Empirical evidence of so-called “selection on the environmental component of the phenotype” (Bonnet et al. 2017) is consequently likely evidence of a genotype-by-environment interaction: something that cannot be detected with widely used statistical analyses of observational data (Kruuk 2004).

## Appendix 2

Four approaches can be used to specify how inheritance alters the distribution of breeding values between selected parents and their offspring. Approach 1: A convolution of the distribution of breeding values in mothers and fathers is taken to construct a distribution of mid-parental values assuming random mating. The distribution of mid-parental values is then convolved with a normal distribution with a mean of 0 and a constant variance to construct the distribution of breeding values in offspring. Approach 2: Breeding values in the offspring generation are assumed to be normally distributed, with a mean equal to that of the mid-point parental breeding value distribution, and a constant variance that does not change across generations. Approach 3: The distribution of breeding values of parents is operated on by an inheritance function which maintains the mid-parental mean value and injects Gaussian distributed segregation variance around each mid-point breeding value. Approach 4: The distribution of breeding values in parents and offspring is identical. As long as selection does not generate distributions with substantial skew or kurtosis, the first three of these approaches give very similar predictions. Various properties of these four approaches are summarised in Table A2.1

**Table A2.1.** Properties of transition functions for breeding values.

|  | Approach 1 | Approach 2 | Approach 3 | Approach 4 |
| --- | --- | --- | --- | --- |
| Are the breeding values in offspring normally distributed? | not necessarily | yes | not necessarily | not necessarily |
| Does the distribution of breeding values in mid-parents and offspring have the same mean? | yes | yes | yes | yes |
| Does the distribution of breeding values in parents and offspring have the same shape? | no | no | no | yes |

## Appendix 3

The Soay sheep model consisted of the following parameterisations:

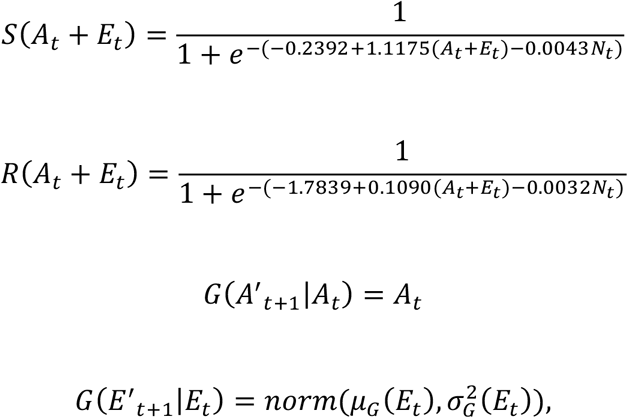

where *μ_G_*(*E_t_*) = 10.603 + 0.6049*E_t_* – 0.0026*N_t_* and 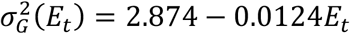

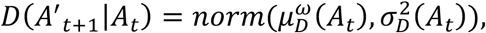

where 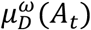 is the mean breeding value post-fertility selection, and 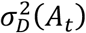 is the additive genetic variance in offspring at birth, which we assumed to be constant and Gaussian distributed. Finally,

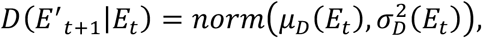

where *μ_D_*(*E_t_*) = 9.8065 + 0.2544*E_t_* – 0.0069*N_t_* and, 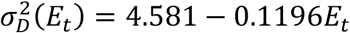.

We started each simulation with mean values of *Ā_t_* = 5.5 and *Ē_t_* = 19.5, an additive genetic variance of 4 and a variance in the environmental component of 6. Because the genetic variances influence the rate of evolution, and selection operate on the entire phenotype, the model results are insensitive to the starting values of *Ā_t_* and *Ē_t_*. Rather it is their sum that determines dynamics. In an initial (unreported) model we tracked a multivariate distribution of age-related breeding values with genetic correlations close to unity. We simplified the model to so each age-related trait has the same breeding value – i.e. genetic correlations are equal to unity. The simplification made no material difference to the results we report.

